# Inducible alpha-synuclein overexpression affects human Neural Stem Cells behavior

**DOI:** 10.1101/290619

**Authors:** J Zasso, M Ahmed, A Cutarelli, L Conti

**Affiliations:** Centre for Integrative Biology – CIBIO, Università degli Studi di Trento, Trento, Italy

## Abstract

Converging evidence suggest that levels of alpha-Synuclein (aSyn) expression play a critical role in Parkinson’s disease (PD). Several mutations of the SNCA gene, encoding for aSyn have been associated to either the familial or the sporadic forms of PD. Nonetheless, the mechanism underlying wild type aSyn-mediated neurotoxicity in neuronal cells as well as its specific driving role in PD pathogenesis has yet to be fully clarified. In this view, the development of proper *in vitro* cellular systems is a crucial step.

Here we present a novel human Tet-on hNSC cell line, in which aSyn timing and level of expression can be tightly experimentally tuned. Induction of aSyn in self-renewing hNSCs leads to progressive formation of aSyn aggregates and impairs their proliferation and cell survival. Furthermore, aSyn induction during the neuronal differentiation process results in impaired neurogenic potential due to enhanced refractoriness to exit self-renewal and to increase of gliogenic *vs* neurogenic competence. Finally, acute aSyn induction in hNSC-derived dopaminergic neuronal cultures results in cell toxicity.

This novel conditional *in vitro* cell model system may be a valuable tool for dissecting of aSyn pathogenic effects in hNSCs and neurons and in developing new potential therapeutic strategies.

## Introduction

Lewy bodies (LBs) are considered a capital hallmark of Parkinson’s Disease (PD). They consist of cytoplasmatic inclusions mainly composed of fibrillar misfolded α-synuclein (aSyn) aggregates within the brain parenchyma [1]. aSyn is a small 140-residue protein highly abundant in neurons and particularly localised at the presynaptic terminals, where it is thought to play roles in vesicles trafficking and in the assembling of the SNARE-complex for neurotransmitters release [2]. Mutations of the *SNCA* gene, encoding for aSyn, are either associated to rare familial PD or represent risk factors for sporadic PD [3, 4]. Furthermore, increased SCNA copy number variation has been shown to be causal for PD, thus suggesting that increased aSyn protein levels are sufficient to trigger the disease [5]. In PD patients, aggregation of aSyn in LBs has been shown, resulting in its mislocalization and loss of function, ultimately leading to the loss of dopaminergic neurons. Beside its role in mature neurons, *in vivo* aSyn overexpression has been shown to affect proliferative potential of mouse hippocampal NSCs, resulting in a reduction of the pool of neural progenitors and decreased neuronal differentiation and maturation [6–9]. Also, ectopic expression of aSyn was shown to affect the migration of NSCs in mouse subventricular zone [10].

Despite the association to PD has been known for decades, aSyn function in human Neural Stem cells (hNSCs) and neurons as well as its involvement in PD is still poor understood. So far, the main limitations have been due to the inaccessibility of adequate *in vitro* cellular models. Indeed, most of the studies aimed at defining a clear role for aSyn in normal and PD neurons have been so far carried out using either non-human systems, including rodents culture [6,7,10], non-neuronal transformed cell lines [11] or non CNS-derived neuronal-like cells [12, 13].

More recently, the advent of hiPSCs technology has widen opportunities to model human conditions, including the possibility to generate *bona fide* human NSCs and mature neurons and consequently matching the affected cells in neurodegenerative disorders [14]. Human hiPSC lines derived from PD patients carrying triplication of the *SNCA locus* have been reported [15]. These cells were proven to efficiently differentiate and mature into TH^+ve^ dopaminergic neurons while showing a two-fold increase in aSyn protein levels with respect to hiPSCs derived from unaffected relatives and recapitulating some aspects of the patient physiology. Interestingly, neuronal precursor cells derived from these PD hiPSCs have been reported to exhibit significant deficiencies in growth, viability, cellular energy metabolism and stress resistance [16], thus indicating that aSyn may play important roles in hNSC physiology. Similarly, wt aSyn overexpression in fetal cortex-derived hNSCs has been shown to impair cell growth and neuronal *vs* glial lineage commitment [17]. Finally, aSyn overexpression has been described to reduce the number of mouse secondary neurospheres formed and to affect NSC morphology and cell cycle progression, leading to their accelerated differentiation [10]. These studies suggest that there is a link between neurogenesis, aSyn and neurodegenerative diseases [18].

Here we report the generation of a novel *in vitro* cell system in which levels and timing of human wt aSyn expression can be experimentally tightly controlled in hiPSC-derived long-term expandable NSCs. Following induction, a progressive increase of aSyn levels can be achieved, leading to the formation of aSyn cytoplasmatic aggregates. This versatile system allows to investigate the effects of aSyn expression on hNSCs behaviour, including self-renewal and differentiation programmes. Our results show that induction of aSyn leads to a reduction in hNSCs growth accompanied by an increased susceptibility to apoptosis. During neuronal differentiation process, aSyn induction affects neurogenic potential by preventing exit from self-renewal and by inducing a shift from neurogenic to gliogenic competence. Further, acute aSyn induction resulted in enhanced apoptotic cell death in hNSC-derived dopaminergic neurons.

This novel *in vitro* cell system may represent a valuable tool for studying aSyn-driven pathogenic-relevant mechanisms in hNSCs and in their mature derivatives and for screening potential therapeutics.

## Materials and methods

### Cell cultures

AF22 cells (kindly donated by Prof. Austin Smith, Cambridge Stem Cell Institute, University of Cambridge, UK) were previously described [19]. Cells were routinely passaged every 2-3 days at a density of 2.5-3.5 x 10^4^ cells/cm^2^ in hNSCs Self-renewal medium composed of DMEM/F-12 (Thermo Fisher Scientific), N2 Supplement (1%, Thermo Fisher Scientific), B27 Supplement (01%, Thermo Fisher Scientific), EGF (10 ng/ml, Peprotech), bFGF (10 ng/ml, Peprotech) and Glutamax (2mM, Thermo Fisher Scientific). Briefly, before seeding the cells, plastic culture vessels were treated with 3 µg/ml Laminin (Thermo Fisher Scientific) at 37 °C for 3-5 hours. For passaging, cells were incubated for 1-2 minutes with StemPro Accutase (Thermo Fisher Scientific) and centrifuged at 260g for 3 minutes. Pellet was resuspended in fresh medium and plated onto the laminin-coated vessels.

For general neuronal differentiation, cells were seeded on laminin-coated cell culture-grade plasticware at a density of 8×10^3^ cells/cm^2^ in self-renewal medium. The following day, medium was shifted to self-renewal medium deprived of EGF and bFGF. After 3 additional days, medium was replaced with neuronal differentiation medium composed of a mix 1:1 of Neurobasal (Thermo Fisher Scientific), and DMEM/F-12, N2 Supplement 1%, B27 Supplement 1%, cAMP 300 ng/ml and Glutamax. For aSyn induction during differentiation process, medium supplemented with doxycycline was replaced every 24h.

For dopaminergic neuronal differentiation, self-renewing cultures were maintained in presence of 200 ng/ml of both FGF8 and SHH for 2-3 weeks to specify a ventral midbrain dopaminergic fate, prior to differentiate them as in the general neuronal differentiation. 750 ng/ml of doxycycline were added at 21 DIV every 24h for 4 DIV before fixing the cells for immunocytochemistry.

### Lentiviral particles preparation and AF22 cells infection

Lentiviral particles carrying the pLVX-TetOne-Puro-human alpha-synuclein vector (kindly donated by Dr. Tilo Kunath, University of Ediburgh, UK) were prepared. This vector, based on the pLVX-TetOn-Puro plasmid (Clontech) is specifically designed to carry on a single vector the cDNA for a Tetracycline transactivator (Tet-on 3G) and a Tetracycline responsive promoter (TREGS promoter) containing 7 Tetracycline Responsive Elements (TRE) controlling the expression of the cDNA for human aSyn. The vector also carries a puromycin resistance cassette for selection of infected cells.

For lentiviral particles preparation, 6×10^4^ HEK 293T cells/cm^2^ were seeded in DMEM containing 10% FBS and left undisturbed overnight. The following day, cells were transfected by CaPO4 method with 20 μg of pLVX-TetOne-Puro-human aSyn, 15 μg of psPAX2 vector (kindly provided by Prof. M. Pizzato, University of Trento, Italy), and 5 μg of VSV-G. Two days after transfection, supernatant was collected and concentrated using the Lenti-X concentrator reagent (Clontech) following the manufacturer recommendation.

For AF22 cells infection, 10 µL of concentrated lentiviral particles were used to infect 2 x 10^4^ cells/cm^2^. After 8h, the medium was completely changed and 72h later, positive selection of the transduced cells was started with 0.3 µg/ml puromycin (Thermo Fisher Scientific) until the non-infected control cells died completely. Cultures were subsequently selected with higher doses of puromycin as discussed in the Results section.

### Cell growth assay

For growth assay, 2 x 10^3^ cells/cm^2^ were seeded onto laminin-coated 24-well plates. aSyn induction was achieved by treatment with 750 ng/ml doxycycline added to the cultures and the medium was renewed every day. Cells were fixed at specific time points by using 4% paraformaldehyde for 15 minutes RT and then nuclei were stained with Hoechst 33258 (Thermo Fisher Scientific). Cell counting was performed using Operetta High-Content Imaging System (Perkin Elmer). Images were collected with a 10x long WD objective considering technical quadruplicate for each time point and the cell number was determined using the software Harmony 4.1 (Perkin Elmer) by the segmentation of the nuclear region.

### Immunocytochemistry and evaluation of aSyn aggregates

For immunofluorescence assay, cells were fixed with 4% paraformaldehyde for 15 minutes at RT, then permeabilised with 0.5% Triton X-100 in PBS for 15 minutes at RT and blocked with blocking solution (5% FBS, 0.3% Triton X-100 in PBS) for 2h at room temperature. Cultures were then incubated O/N at 4°C with specific primary antibodies (see Table S1) diluted in blocking solution. After three rinses with PBS, cells were incubated with the appropriate secondary antibodies (see Table S1) for 2 hours and nuclei counterstained with Hoechst 33258 before imaging with a Leica DMIL inverted fluorescent microscope. For the quantification of specific immunopositive cells, at least 3000 cells per condition for every antigen were counted. Data were normalised on the total number of cells in every field.

For evaluation of aSyn aggregation, cells were treated with 750 ng/ml of doxycycline for 12 days. Cultures were fixed with 4% paraformaldehyde for 15 minutes at RT and stained with anti-*α*-synuclein antibody. Nuclei and cytoplasm were counterstained using Hoechst 33258 and CellMask Deep Red (Molecular Probes). Cells were analysed using the Operetta High-Content Imaging System. Images were collected using a 20x long WD objective considering technical quadruplicate and analysed with Harmony 4.1 software. Briefly cells were identified by the segmentation of the nuclear region based on the Hoechst 33258 signal and cytoplasm region of interest (ROI) was defined based on the CellMask signal. aSyn aggregates were detected as fluorescent spots inside the ROI. Number of objects, area and intensity of the signal were calculated for each well and normalised on the number of aSyn^low/Med^ and aSyn^high^ intensity aSyn immunoreactive cells.

### Western Blot analysis

Cells were lysed using the SDS sample buffer (62.5 mM Tris-HCl pH 6.8, 2% SDS, 10% Glycerol, 50 mM DTT) and boiled for 5 minutes at 95°C before loading into a 15% polyacrylamide gel run at 15 mA. After transfer on PVDF membrane at 100V constant for 2hrs, proteins were incubated in 0.4% paraformaldehyde in phosphate buffered saline solution for 30 min at RT as previously reported [20] and then in blocking solution (10% milk) in TBS-T for 1 hr. Membranes were further incubated O/N at 4°C in agitation with primary antibody. After the washing step with TBS-T, membranes were incubated with secondary antibody for 2 hrs. Both primary and secondary antibodies were prepared in TBS-T supplemented with 5% non-fat milk diluted in TBS-T. Signal was detected using Clarity ECL reagents (Biorad) in a dark chamber Uvitec Alliance (Uvitec) and the manufacturer software to acquire and analise the data.

### Statistical Analysis

Statistical significance was assayed using a Student’s unpaired t test or two-way ANOVA analysis of variance using the GraphPad Prism software. P < 0.05 was considered statistically significant.

## Results

### 1. Modulation of wt aSyn expression in AF22 Tet-On aSyn cells

To generate a hNSC line for the inducible expression of wt aSyn, hiPSC-derived AF22 cells were infected with lentiviral particles carrying the pLVX-TetOn-Puro-human *α*-Syn vector. Following infection, cultures were exposed to 1 μg/mL puromycin, chosen as optimal puromycin dose that allows for effective selection without altering the normal self-renewal potential of the resistant cells (Fig. S1A). AF22 Tet-On aSyn cells selected with this puromycin dose exhibited low aSyn basal expression levels that increased 7.7-fold following treatment with doxycycline for 72 hrs (Fig. S1B). Based on our previous experience with Tet-On systems, the abovementioned induction experiments were performed using a 750 ng/mL dose of doxycycline. To test if AF22 Tet-On aSyn cells show a dose-dependent level of induction and to test at which dose of doxycycline it is reached the maximum level of induction, we treated the cultures for 72 hrs with different concentrations of doxycycline (0, 100, 250, 500, 750 and 1000 ng/mL). A clear dose-response relationship in the induction of aSyn expression levels was found, reaching a maximum plateau of 8.0-fold induction already at 750 ng/mL doxycycline treatment (Fig. S2A). Higher doses of doxycycline did not lead to significative increase of aSyn induction levels. Based on these results, we confirmed 750 ng/mL doxycycline as optimal dose for the further analyses reported in this study.

Short-term time course analysis of doxycycline induction on AF22 Tet-On aSyn cells (Fig. 1A) showed a progressive increase of aSyn expression levels along with time of doxycycline treatment (2.2-, 5.3-and 7.9-fold induction at 24, 48 and 72 hrs of doxycycline treatment, respectively; Fig. 1B). Immunofluorescence assay performed on these cultures showed that not all of the aSyn immunoreactive cells exhibited the same levels of expression, with even a fraction of the cells that barely showed any transgene expression (Fig. 1C). Quantitative analysis revealed that 72.21±16.64 % of cells in culture were aSyn^+ve^, the remaining showing very low or undetectable aSyn immunoreactivity (Fig. 1D). Further, the pool of aSyn^+ve^ cells can be divided in 30.3±8.21 % cells exhibiting high aSyn immunoreactivity (aSyn^high^ cells) and 69.67±8.05 % cells exhibiting low/medium aSyn immunoreactivity (aSyn^low/Med^ cells) (Fig. 1E).

**Figure 1.**
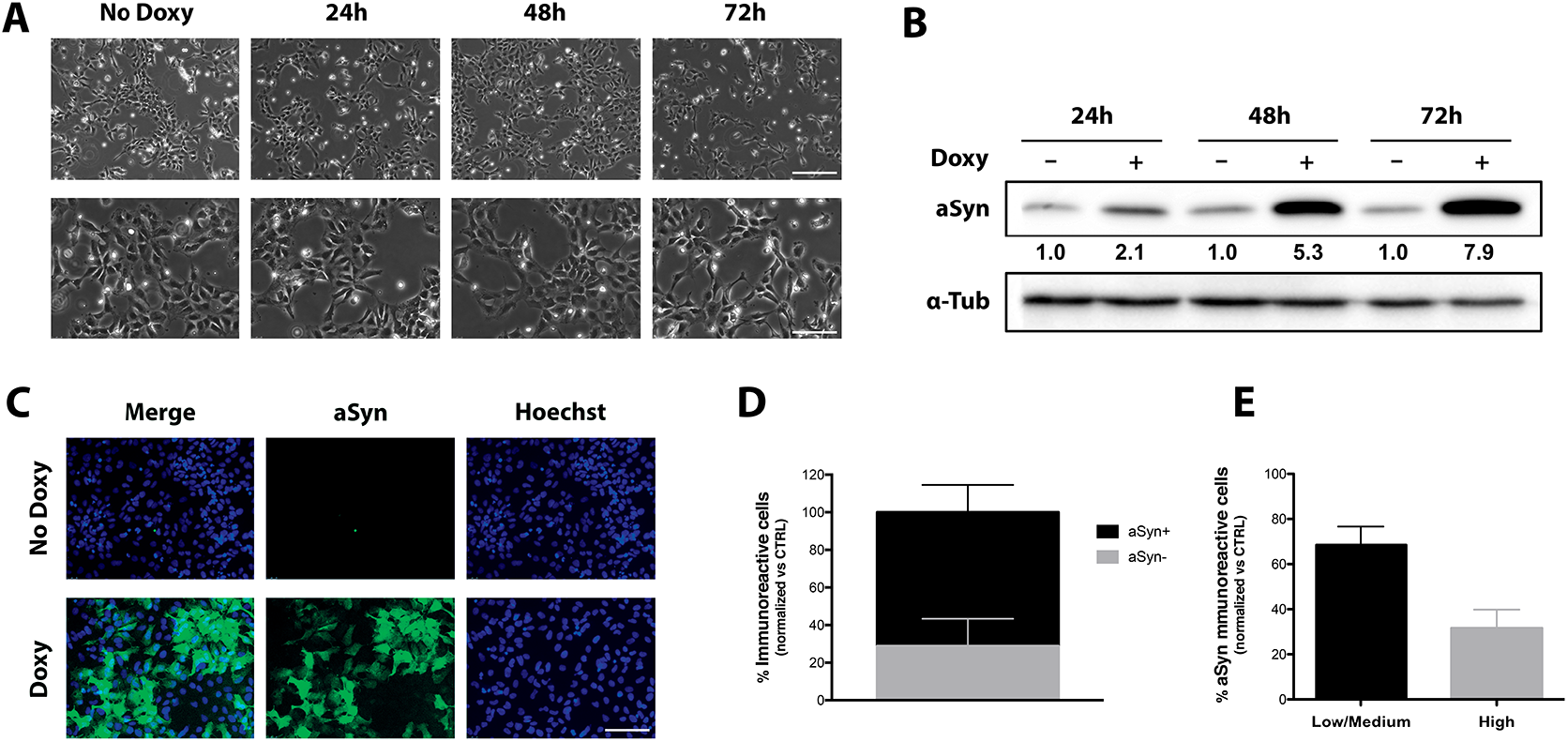
Modulation of wt aSyn expression in AF22 Tet-On aSyn cells. **A-B.** Time course of αSyn induction in AF22 Tet-On aSyn cells treated with 750 ng/ml doxycycline. **A.** Phase contrast pictures of AF22 Tet-On aSyn cells during time course induction of αSyn. Scale bars: upper panels 200 μm; lower panels 100 μm. **B.** Representative image of western blot assays of time course induction of aSyn. Densitometric quantification was normalised on α-tubulin expression and versus untreated cells. **C-E.** Immunocytochemistry analysis for αSyn on AF22 Tet-On aSyn cells untreated and treated with 750 ng/ml doxycycline for 72h and quantification of immunopositive cells. **C.** Representative immunofluorescent images showing αSyn^+ve^ cells in AF22 Tet-On aSyn cultures. Hoechst was used for nuclear staining. Scale bar: 100 μm. **D.** Quantification of the total number of αSyn^+ve^ cells in basal and induced conditions. **E.** Quantification of the percentage of aSyn^high^ cells and aSyn^low/Med^ cells among the overall population of αSyn^+ve^ cells in induced AF22 Tet-On aSyn cells.

Longer time course analysis of doxycycline induction on AF22 Tet-On aSyn cells (Fig. S2B) confirmed the progressive increase of transgene expression levels from 48 to 72 hrs of doxycycline induction and a strong immunoreactive band at 12 days of induction (2.0-, 7.4-and 13.4-fold induction at 48 and 72 hrs and 12 days of doxycycline treatment, respectively; Fig. S2C). On the whole, these results indicate that aSyn induction in AF22 Tet-On aSyn cells is long-term maintained and enlightened progressive aSyn accumulation.

### 2. wt aSyn over-expression affects cell division and cell viability in AF22 Tet-On aSyn cells

To test if aSyn induction in the cultures results in phenotypic abnormalities on hNSCs growth, we performed a cell growth assay on AF22 Tet-On aSyn cells. Since aSyn has been described to influence mitochondrial activity, we decided to avoid mitochondrial activity-based growth assays. We thus performed an automated HTS cell count of stained cell nuclei on cultures fixed at defined time points.

aSyn induction produced a slight reduction in cell growth already after 72 hrs and this effect was more marked at later time-points (Fig. 1A and Fig. 2A), where a strong impairment in the growth occurred (% reduction in cell number: 34.12 and 44.74 at 5 and 7 days, respectively; Fig. 2A). These data could be interpreted by possible effects elicited by aSyn either on (*i*) cell division, (*ii*) cell death, (*iii*) change of fate by induction of differentiation or (*iv*) a combination of the abovementioned effects. In order to dissect out which of these possibilities was prominent, we performed specific assays. Analysis of phospho-Histone H3^+ve^ cells present in the cultures at different time points (Fig. 2B) showed a reduction, although not statistically significative, of phospho-Histone H3^+ve^ cells occurring both at 72 and 96 hrs of induction (Fig. 3C). A statistically significant 47.16% reduction was appreciated at 120 hrs (Fig. 3C). Further, a stronger statistically significative reduction of phospho-Histone H3^+ve^ cells in aSyn^+ve^ cells compared to aSyn^-ve^ cells, at all the time points considered (% of reduction of phospho-Histone H3^+ve^ cells in aSyn^+ve^ cells in culture: 82.3, 96.79 and 91.54 at 3, 4 and 5 days, respectively) (Fig. 3D). These results indicate that aSyn induction affects cell division in AF22 Tet-On aSyn cells. Further, immunofluorescent analysis for NSC markers (Nestin and sox2), neurons (*β*3-Tubulin) and astrocytes (GFAP) indicated that more than 97% of the cells retain their normal NSC identity without any significative induction of neuronal or glial cells (not shown), thus indicating that aSyn overexpression in self-renewing conditions doesn’t force differentiation.

**Figure 2.**
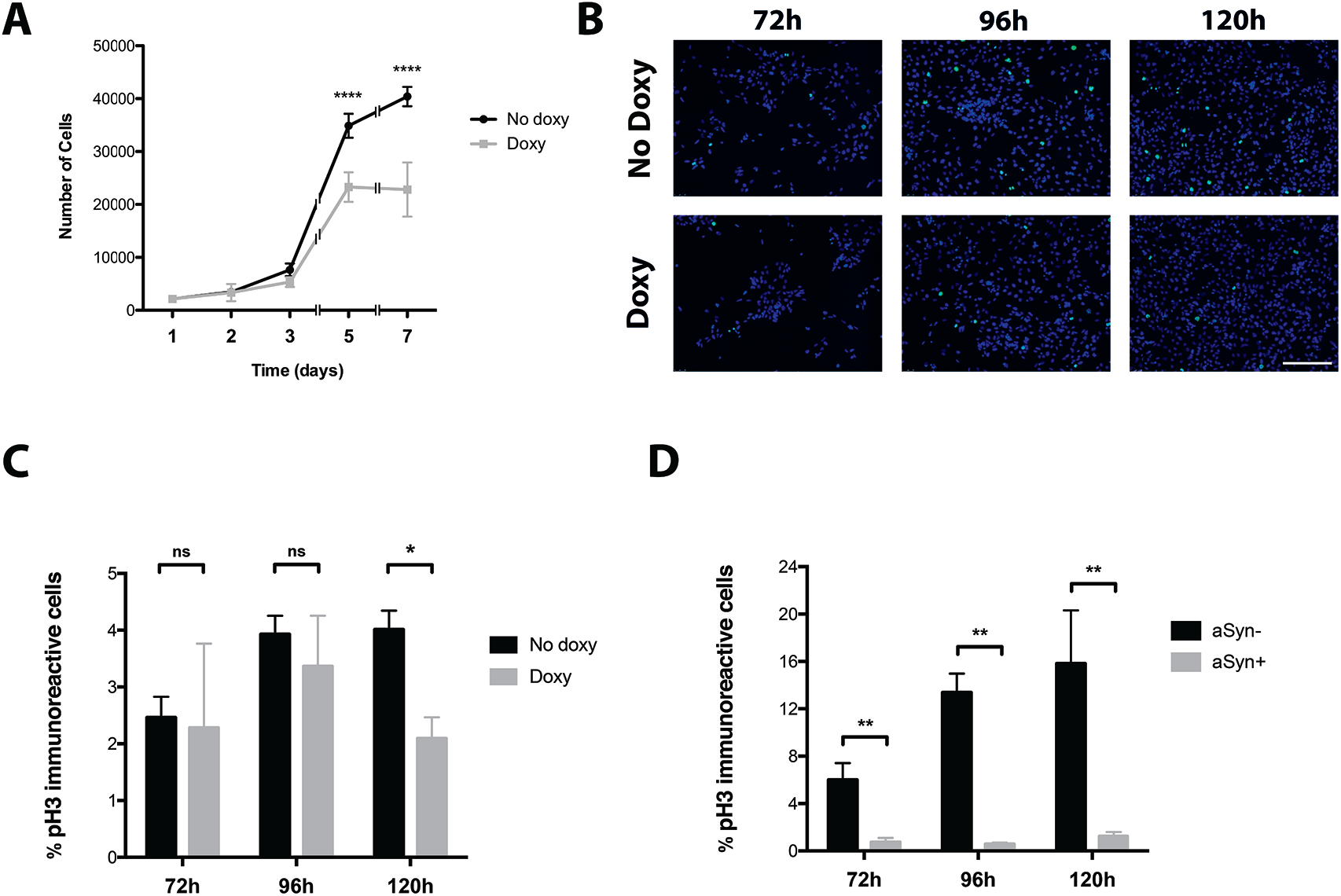
wt αSyn expression affects proliferation of AF22 Tet-On aSyn cells. **A.** Cell growth assay on basal or induced AF22 Tet-On aSyn cells. **B-D.** Time course immunofluorescent analysis for phospho-Histone H3 on AF22 Tet-On aSyn cells untreated or treated with 750 ng/ml doxycycline and relative quantification of immunopositive cells. **B.** Representative pictures of AF22 Tet-On aSyn cells stained for pospho-Histone H3. Hoechst was used for nuclear staining. Scale bar: 200 μm. **C.** Quantification of the total number of pospho-Histone H3 immunopositive cells at defined time points. **D** Quantification of the percentage of pospho-Histone H3 immunopositive cells at defined time points in induced AF22 Tet-On aSyn cells. Normalization was performed on the populations of αSyn^+ve^ and αSyn^-ve^ cells. *ns*: non significative, *: p < 0.05, **: p < 0.01.

**Figure 3.**
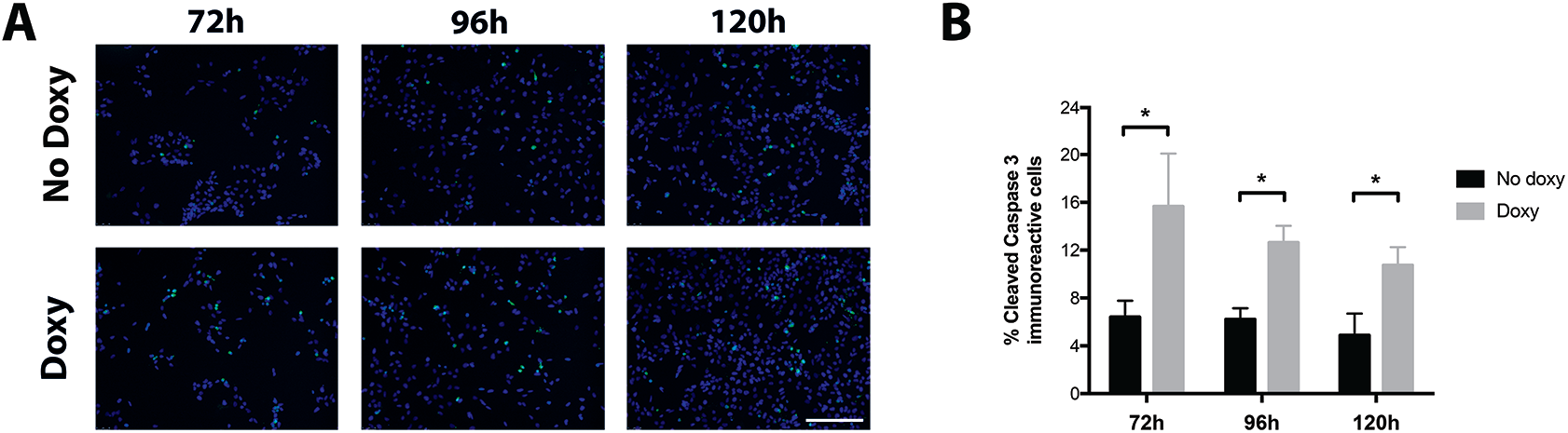
wt aSyn over-expression affects cell viability in AF22 Tet-On aSyn cells. **A-B.** Immunocytochemistry analysis for cleaved Caspase 3 on AF22 Tet-On aSyn cells treated or not with 750 ng/ml doxycycline and relative quantification of immunopositive cells. **A.** Representative pictures of uninduced or induced AF22 Tet-On aSyn cells stained for cleaved Caspase 3. Hoechst was used for nuclear staining. Scale bar: 200 μm. **B.** Quantification of the cleaved Caspase 3^+ve^ cells at the different time points. Number of cleaved Caspase 3^+ve^ cells was normalised on the total number of cells. *: p < 0.05.

In order to test if aSyn induction could also impact on cell survival, we performed an immunofluorescent analysis for cleaved Caspase-3 at different time points following doxycycline treatment (Fig. 3A). aSyn induction led to a marked increase in the number of apoptotic cells in culture (fold increase of cleaved Caspase-3^+ve^ cells in culture: 3.31, 2.03 and 2.19 at 3, 4 and 5 days, respectively) (Fig. 3B).

On the whole, these results indicate that aSyn overexpression in hNSCs leads to a reduced cell growth by a double action, both by affecting proliferation capability and by enhancing cell death occurrence.

### 3. Conditional overexpression of wt aSyn leads to formation of intracellular aggregates in AF22 Tet-On aSyn cells

aSyn has been shown to generate aggregates inside of the cells and that these may contribute to cellular dysfunctions. Immunofluorescence analysis for aSyn in AF22 Tet-On aSyn cells following 12 days of doxycycline treatment showed that a fraction of aSyn^+ve^ cells exhibited punctate immunoreactive dots in the cytoplasm (arrows in Fig. 4A). aSyn aggregates were present both in aSyn^high^ cells and aSyn^low/Med^ cells, although the former presented higher number of aggregates with respect to the latter (number of aggregates: 0.48±0.19 and 3.14±0.32 in aSyn^low/Med^ and aSyn^high^ cells, respectively; Fig. 4B). Also, aggregates in aSyn^high^ cells exhibited a 5.8-fold increase in the immunoreactive signal with respect to the ones present in aSyn^mod/low^ cells (Fig. 4C). Also, aSyn^high^ cells showed an 3.1-fold increase in the parameter of area of aggregated spots with respect to aSyn^mod/low^ cells (Fig. 4D).

**Figure 4.**
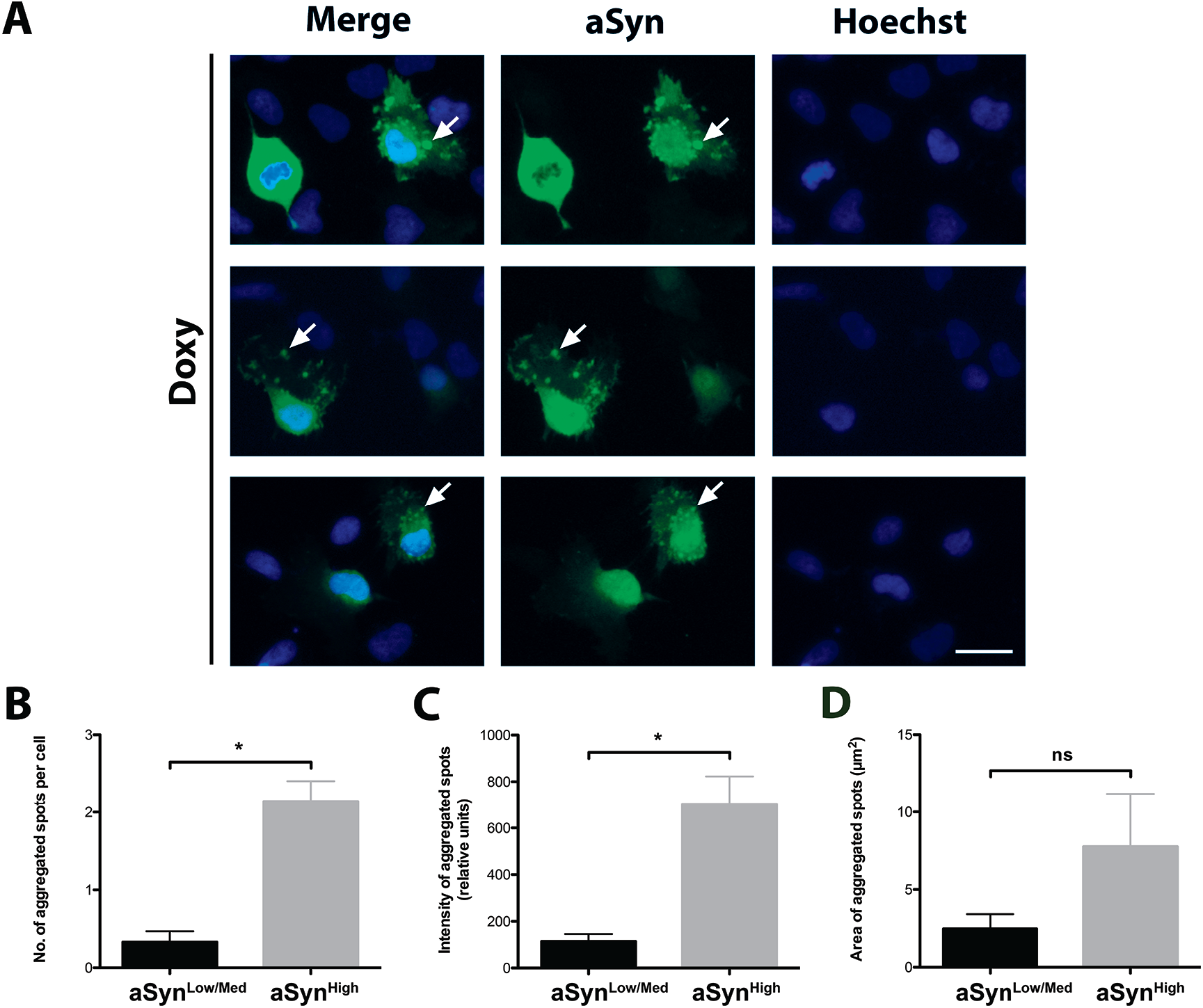
Conditional overexpression of wt aSyn leads to formation of intracellular aggregates in AF22 Tet-On aSyn cells. **A-D.** Immunocytochemistry analysis for αSyn on AF22 Tet-On aSyn cells treated with 750 ng/ml doxycycline for 12 days and relative evaluation of aSyn aggregates. **A.** Representative pictures of AF22 αSyn Tet-On treated with 750 ng/ml doxycycline for 12 days and stained for aSyn and Hoechst. Arrows indicate αSyn aggregates. Scale bar: 25 μm. **B.** Quantification of the number of αSyn aggregates per cell in aSyn^Low/Medium^ and αSyn^High^ cells. **C.** Quantification of the intensity of the fluorescent signal of the αSyn aggregated spots in aSyn^Low/Medium^ and αSyn^High^ expressing cells. **D.** Quantification of the area of aSyn aggregated spots in aSyn^Low/Medium^ and αSyn^High^ expressing cells. *ns*: non significative, *: p < 0.05.

These results indicate that aSyn over-expression produces aggregation on long-term induced AF22 Tet-On aSyn cells and that the occurrence of aggregation is dependent on the level of aSyn over-expression, with aSyn^high^ cells producing more aggregated spots with greater intensity with respect to aSyn^mod/low^ cells.

### 4. aSyn overexpression impairs neuronal differentiation in AF22 Tet-On aSyn cells

aSyn overexpression has been indicated as a main player in neuronal dysfunction. To test if aSyn induction in AF22 Tet-On aSyn cells could induce defects in the neuronal differentiation potential of the cultures, AF22 Tet-On aSyn cells (CTRL and cells maintained in doxycycline for the entire differentiation procedure) were exposed to a 2-week neuronal differentiation protocol. In these conditions, cell started to show morphological changes indicative for their progressive neuronal maturation. Immunofluorescence analysis on 14 days induced cultures showed that 73.25±1.97 % of the cells in culture were aSyn^+ve^ (not shown). As expected, at this stage, not induced cultures were mainly composed of *β*3-tubulin^+ve^ neurons (% of *β*3-tubulin^+ve^ cells: 87.52±6.09; Fig. 5A and C) with only a fraction of the cells in culture positive for Map2, a marker for mature neurons (% of Map2^+ve^ cells: 21.14±4.35; Fig. 5B and C). On the contrary, doxycycline-treated cultures showed a 51.85% and 38.45% reduction in the overall number of *β*3-tubulin^+ve^ cells and Map2^+ve^ cells, respectively (Fig. 5A-B-C). These results indicate that aSyn overexpression partially affects neuronal differentiation capability of hNSCs. We next asked if the reduction in the number of neuronal cells is mainly due to (*i*) an impaired competence of the cells to start the neuronal differentiation process or (*ii*) induced competence to differentiate toward non-neuronal fates (i.e. shift from neurogenic *vs* gliogenic fate). Sox2 immunofluorescence (Figure S3) showed the presence of large clusters of sox2^+ve^ cells in the doxycycline-treated culture with a 12-fold increase in the number of sox2^+ve^ cells thus indicating that aSyn overexpression increased the percentage of the cells refractory to undergo neuronal differentiation process (Fig. 5D). Furthermore, aSyn overexpression induced a 7.61-fold increase in the number of GFAP^+ve^ cells (% of GFAP^+ve^ cells: 0.46±0.23 and 3.57±0.61 in not induced and doxycycline-induced cells, respectively; Fig. 5D) in the cultures, thus indicating an enhanced propensity of the cells to differentiate toward the astrocytic lineage.

**Figure 5.**
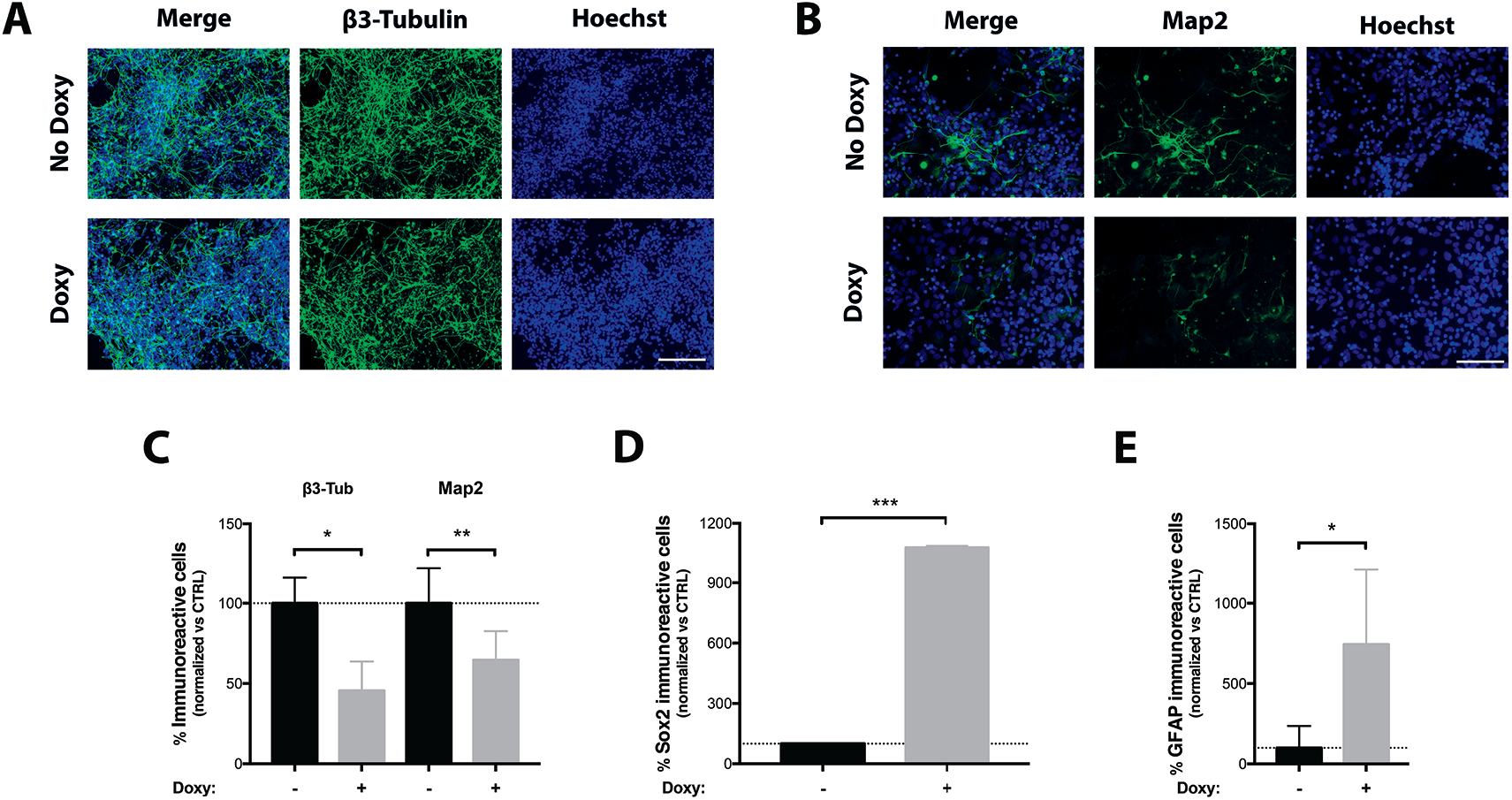
aSyn expression impairs neuronal differentiation of AF22 Tet-On aSyn cells. **A-E.** Effects of aSyn expression on neuronal differentiation of AF22 Tet-On aSyn cells treated or not with 750 ng/ml of doxycycline for the entire differentiation process (14 DIV). **A.** Representative pictures of β3-Tubulin^+ve^ cells on 14 DIV cultures of basal and induced AF22 Tet-On aSyn cultures. Scale bar: 100 μm. **B.** Representative pictures of Map2^+ve^ cells on 14 DIV cultures of basal and induced AF22 Tet-On aSyn cultures. Hoechst was used for nuclear staining. Scale bar: 100 μm. **C.** Quantification of the total number of β3-Tubulin^+ve^ and Map2^+ve^ cells on basal and induced conditions. Values are normalized over the basal condition. **E.** Quantification of the total number of sox2^+ve^ cells on basal and induced conditions. Values are normalized over the basal condition. **F.** Quantification of the total number of GFAP^+ve^ cells on basal and induced conditions. Values are normalized over the basal condition. *: p < 0.05, ***: p < 0.01, ***: p < 0.0001.

On the whole, these results indicate that there is a combined effect of aSyn over-expression in disturbing hNSC neuronal differentiation process both by decreasing the competence of the cells to respond to neuronal differentiative environment and by shifting neurogenic to gliogenic fate.

### 5. aSyn overexpression affects cell viability in AF22 Tet-On aSyn cell-derived dopaminergic neurons

AF22 cells have the competence to differentiate toward dopaminergic neuronal fate in defined *in vitro* conditions [19]. Thus, we next analysed the possible effects elicited by aSyn acute induction in long-term differentiated hNSC-derived dopaminergic neuronal cultures. To this aim, self-renewing AF22 Tet-On aSyn cells were patterned for three weeks with FGF8/SHH and then induced to differentiate to dopaminergic neurons for three weeks as previously reported [19, 21]. After three weeks of dopaminergic neuronal maturation, cultures were mainly composed of neurons (% of *β*3-tubulin^+ve^ cells: 86.53±7.17), most of which were dopaminergic Nurr-1^+ve^ neurons (% of Nurr-1^+ve^/*β*3-tubulin^+ve^ neurons: 74,43±8.26 %). At this stage, acute aSyn overexpression was achieved by treating the cultures for 4 days with 750 ng/mL doxycycline. aSyn overexpression produced an acute toxic effect resulting in degeneration of the neuronal cells in culture (% reduction of the total number of neurons in aSyn induced cultures: 24.72±6.13) (Fig. 6A) and by a 1.94-fold increase in the number of cleaved Caspase-3^+ve^ cells (Fig. 6B). Quantitative analysis showed that the fraction of Nurr-1^+ve^/*β*3-tubulin^+ve^ neurons was significantly decreased following aSyn induction (% reduction of the number of Nurr-1^+ve^/*β*3-tubulin^+ve^ neurons in aSyn induced cultures: 31.72±4.36) (Fig. 6C) with only a minor effect on the Nurr-1^-ve^/*β*3-tubulin^+ve^ neurons, that were not affected by acute aSyn overexpression (not shown). Thus, aSyn overexpression impairs the survival of hNSC-derived neurons, with dopaminergic neurons being differentially affected with respect to non-dopaminergic neuronal subtypes.

**Figure 6.**
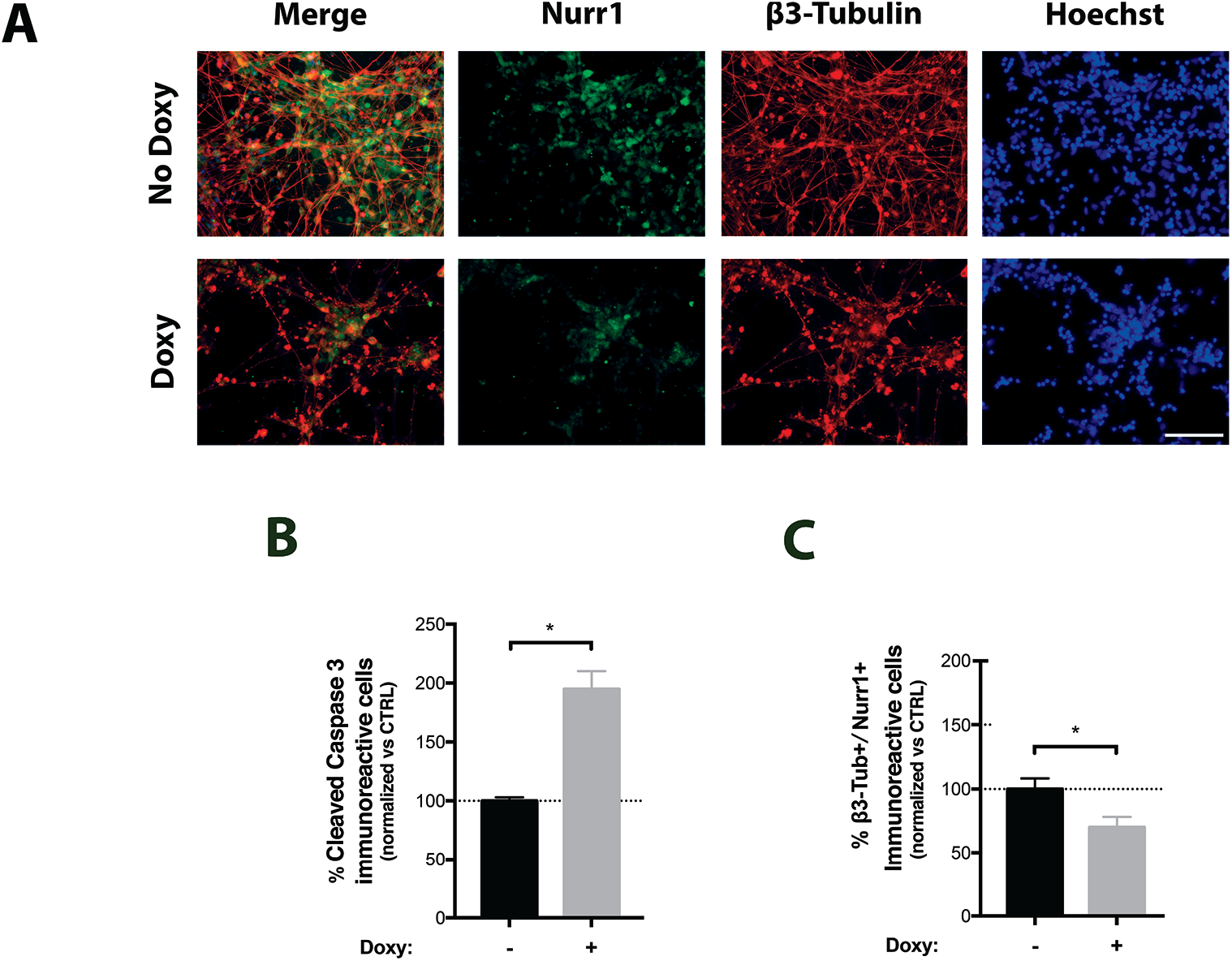
aSyn overexpression affects cell viability in AF22 Tet-On aSyn cell-derived dopaminergic neurons. **A-C.** Effects of acute aSyn induction (21 to 25 DIV) on AF22 Tet-On aSyn cell-derived dopaminergic neurons. **A.** Representative pictures of β3-Tubulin^+ve^/Nurr1^+ve^ cells in 25 DIV cultures. Hoechst was used for nuclear staining. Scale bar: 100 μm. **B.** Quantification of the total number of Cleaved Caspase 3^+ve^ cells on basal and induced conditions. Values are normalized over the basal condition. **C.** Quantification of the total number of β3-Tubulin^+ve^/Nurr1^+ve^ cells on basal and induced conditions. Values are normalized over the basal condition. *ns*: non significative, *: p < 0.05.

## Discussion

Although aSyn is considered of relevance for PD pathogenic process, the molecular mechanism triggering PD starting from aSyn homeostasis alteration is still a matter of debate. Here we report the generation of a novel cellular model based on the overexpression of wt human aSyn in NSCs. The system is characterized by the controlled expression of wt human aSyn by means of a Tet-On inducible mechanism and represents a valuable tool to study the effects of aSyn overexpression in hNSCs and neurons.

The AF22 Tet-On aSyn cell model here developed has several features that make it attractive for studying the biological and molecular effects elicited by aSyn in different developmental paradigms. These include the human NSC nature of the parental AF22 cells. Originally described by Falk and colleagues [19], these cells have been obtained from normal human iPSCs and show features that make them ideal as parental cells to be engineered. Indeed, they are homogeneously composed of self-renewing NSCs characterized by genomic stability and highly amenability to genetic manipulation. Importantly, these cells maintain a stable high neurogenic capability along long-term *in vitro* expansion and the competence to respond to specific patterning cues that allows generating defined neuronal subtype populations, including dopaminergic neurons.

The inducible nature of aSyn expression coupled to the NSC system opens to the possibility to study both acute or chronic aSyn-mediated effects in defined relevant cell populations, i.e. neural progenitors and mature neurons. Also, this is instrumental to sort out specific effects in several processes, including self-renewing, lineage commitment and neuronal maturation and/or maintenance.

aSyn inducible systems have been reported from different immature parental cell lines, mainly PC12 cells and human neuroblastoma lines [12,13,22–24]. Nonetheless, these systems have some intrinsic limitations that are overcome in AF22 Tet-On aSyn cell model. Indeed, PC12 cells are of rodent origin and species-specific differences in aSyn sequence and roles have been reported [25–27]. Additionally, both PC12 cells and neuroblastoma lines are transformed and share a non-CNS origin. Noteworthy, their neurogenic potential is limited in terms of efficiency and quality of neuronal-like cells that can be obtained following their differentiation, thus limiting their physiological relevance for studies aimed at dissecting aSyn roles in human CNS neurons.

AF22 Tet-On aSyn cells show a clear dose-response transgene expression with a robust aSyn induction up to 7-8 fold; also, induction can be longterm maintained leading to a progressive increase in aSyn levels. When we tried to induce aSyn aggregation in AF22 Tet-On aSyn cells, we found that this process requires a prolonged induction period allowing to aSyn levels to increase progressively. The appearance of aSyn aggregates in AF22 Tet-On aSyn cells occurs after 12 days of induction. Other studies performed on PC12/TetOn aSyn inducible systems failed to observe wt aSyn aggregates in proliferating cells [22]. This could be due to the different origin of the parental cells or to the fact that levels of aSyn induction are lower or to the quite reduced time of induction with respect to our study.

It is interesting to note that wt aSyn overexpression induces phenotypic defects in hNSCs in the self-renewing state. Other studies have reported that overexpression of aSyn in the proliferating state fails to induce any cell death in proliferating neural-like cells, despite the prominent accumulation of aSyn aggregates [12]. The factors accounting for the differential death effects could include differences in clearance mechanisms, or involvement of cell cycle molecules or other proteins differentially expressed in the two states.

An increasing number of studies reveal that aSyn may play an important role in neurogenesis. When the SNCA gene is differentially expressed or bears mutations, the *in vivo* NSC pool is negatively regulated and both neurogenesis and survival of newly generated neurons are decreased [9]. These studies suggest that might exist a link between neurogenesis, aSyn and neurodegenerative diseases [18].

*In vitro*, aSyn overexpression has been described to reduce the number of mouse secondary neurospheres formed and to affect NSC morphology and cell cycle progression, leading to their accelerated differentiation [10].

hiPSC-derived neuronal precursor cells from a PD patient carrying a genomic triplication of the SNCA gene showed substantial impairments in growth, viability, cellular energy metabolism and stress resistance. These effects were exacerbated when the cultures were challenged by starvation or toxic stimuli [16]. Also, overexpression of wt aSyn in expanded populations of progenitors derived from the human fetal cortex showed a slight effect on cell growth and a progressive impairment of lineage commitment competence [17]. Similarly, studies on hESC-derived neural progenitors overexpressing wt aSyn and on neural progenitors obtained from hiPSCs fom a PD patient with a SCNA locus triplication, showed an increased cell death and reduced neurogenic capacity compared with control cultures [17, 28].

Our results confirm that overexpression of human wt aSyn impairs the process of transition of human NSCs to neuronal cells. In particular, we have seen a major effect on the efficiency of the cells to exit from self-renewal when exposed to neuronal differentiative cues. It is not yet clear which molecular mechanisms trigger this specific defect. PSC-derived long-term expandable hNSCs, including the AF 22 cells, are highly responsive to efficiently undergo neuronal differentiation when exposed to the neuronal differentiative protocol here employed [19, 21]. Additional investigation is required to define if this defect is maintained when exposing the cultures to other pro-neuronal differentiative conditions. Further, exposure to non-neuronal (i.e. gliogenic) differentiation cues could help to understand if the observed refractoriness to exit self-renewal is specific for the transition toward the neuronal lineage or is a more general aSyn-mediated effect. Beside this, we have here reported a partial shift of competence of the cells from a neurogenic to a gliogenic fate, that results in increased number of astrocytes found in neuronally differentiating cultures. Other studies have reported that wt aSyn overexpression in human neural progenitors derived from fetal cortex preserved the neurogenic competence of the cells following long-term expansion [17]. To this respect, we can speculate that this discrepancy might be related to the different nature and identity between our and the abovementioned cell system. Indeed, differently from fetal brain-derived cell system that is representative of late developmental neural stages and in which the neurogenic competence quickly declines with *in vitro* passages, the PSC-derived hNSCs we employed are representative of earlier developmental neural stages and are extremely stable also following extensive long-term expansion [29].

Finally, we have observed an aSyn-mediated acute toxicity in hNSC-derived neurons, being dopaminergic neurons predominantly affected. These results are in agreement with a previous study reporting acute aSyn toxicity in hESC-derived neuronal cultures [17]. These authors showed that hESC-derived neuronal cultures are highly vulnerable to expression of both wt aSyn or mutant aSyn forms, with dopaminergic neurons exhibiting higher toxic susceptibility with respect to non-dopaminergic (GABAergic) neurons. It is yet unclear the reason of this neuronal sub-types selective cytotoxicity. The factors accounting for this differential death effects in different neuronal subtypes are unknown, but could include differences in clearance mechanisms or other proteins differentially expressed in the two states. Interestingly, aSyn over-expression has been shown to directly affect TH expression, suggesting possible direct TH effects [30]. Regardless of the exact reason, this fact further validates the current model as one in which toxic effects occur preferentially in dopaminergic neurons.

In conclusion, we have developed a cell system for controlled expression of wt aSyn in hNSCs that exhibit defined aSyn-driven phenotypes both in self-renewal and differentiating/differentiated stages. This novel inducible model may prove valuable in the deciphering of aSyn-mediated pathogenic effects and in the assessment and screening of potential therapeutic strategies.

## Acknowledgements

The authors wish to thank Prof. Austin Smith (Cambridge Stem Cell Institute, University of Cambridge, UK) for providing the AF22 cell line, Dr. Tilo Kunath (MRC Centre for Regenerative Medicine, University of Edinburgh, UK) for the pLVX-TetOn-Puro-human alpa-synuclein vector and Prof. Massimo Pizzato for the psPax2 vector. We also thank personnel of Cibio HTS Core Facility and of Advanced Imaging Core Facility. This work was supported by the intramural funding from the University of Trento.

