## Supplementary Materials for "Inducible alpha-synuclein overexpression affects human Neural Stem Cells behavior"

### Supplemental Material

#### Supplemental Figure Legends

**Figure S1. Evaluation of aSyn induction in AF22 Tet-On aSyn cultures selected by means of different doses of puromycin. Upper Panel.** Phase contrast images of AF22 Tet-On aSyn cells selected with different doses of puromycin and treated or not with 750 ng/ml of doxycycline for 72h. Scale bar: 100 µm. **Lower Panel.** Representative image of a western blot analysis of the cells in (A). Densitometric quantification was normalised on α-tubulin expression and versus untreated cells.

**Figure S2. Evaluation of dose-response and long-term aSyn induction in AF22 Tet-On aSyn cultures. A.** Dose-dependent αSyn induction in AF22 Tet-On aSyn cells treated with or without doxycycline for 72h and relative densitometric quantification normalised on α-tubulin expression and versus untreated control. **B.** Phase contrast pictures of AF22 Tet-On aSyn cultures treated with or without 750 ng/ml doxycycline for the indicated time. Scale bar: 100 µm. **C.** Kinetics of αSyn induction of AF22 Tet-On aSyn cells treated with 750 ng/ml doxycycline for the indicated time showing a time dependent accumulation of αSyn levels. Expression is maintained in long-term induced (12 DIV) cultures .

**Figure S3. aSyn induction reduced neurogenesis in in AF22 Tet-On aSyn cells.** Effects of aSyn expression on neuronal differentiation of AF22 Tet-On aSyn cells treated or not with 750 ng/ml of doxycycline for the entire differentiation process (14 DIV). Representative pictures of sox2$^{+ve}$ cells on DIV14 cultures of basal and induced AF22 Tet-On aSyn cultures. Presence of clusters of sox2$^{+ve}$ cells are visible in induced cultures. Hoechst was used for nuclear staining. Scale bar: 100 µm.

**Figure S4. Acute aSyn induction impairs AF22 Tet-On aSyn cell-derived dopaminergic neurons viability.** Effects of acute aSyn induction (21 to 25 DIV) on AF22 Tet-On aSyn cell-derived dopaminergic neurons. Representative pictures of Cleaved Caspase 3$^{+ve}$ and aSyn$^{+ve}$ cells in 25 DIV cultures. Hoechst was used for nuclear staining. Scale bar: 100 µm.

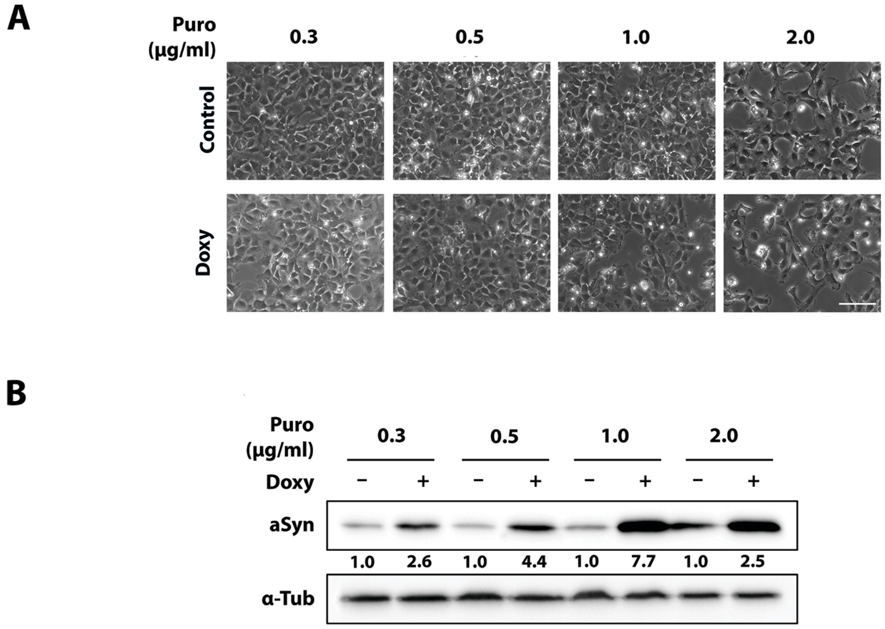


**Figure S1.** Evaluation of aSyn induction in AF22 Tet-On aSyn cultures selected by means of different doses of puromycin.

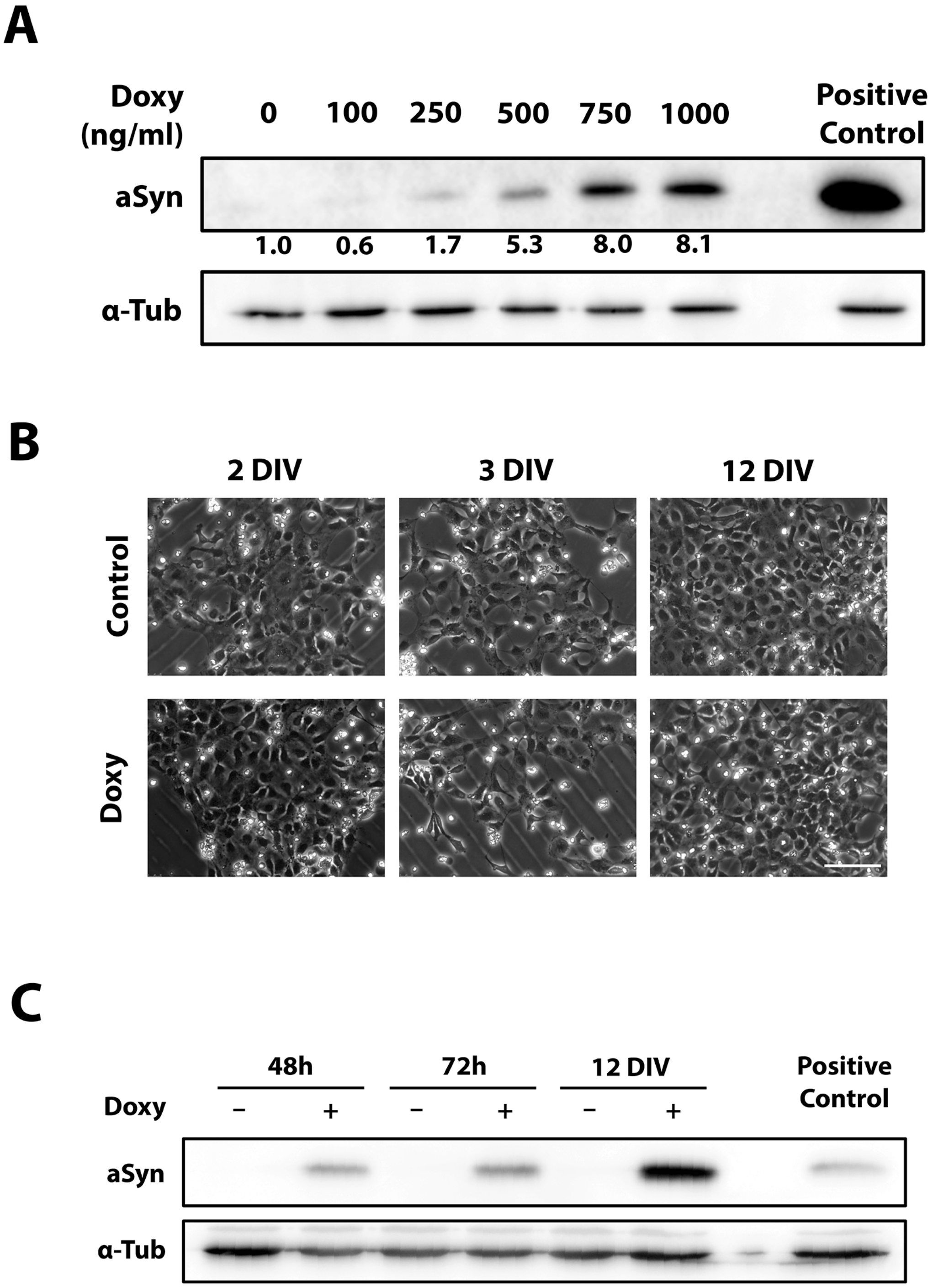


**Figure S2.** Evaluation of dose-response and long-term aSyn induction in AF22 Tet-On aSyn cultures.

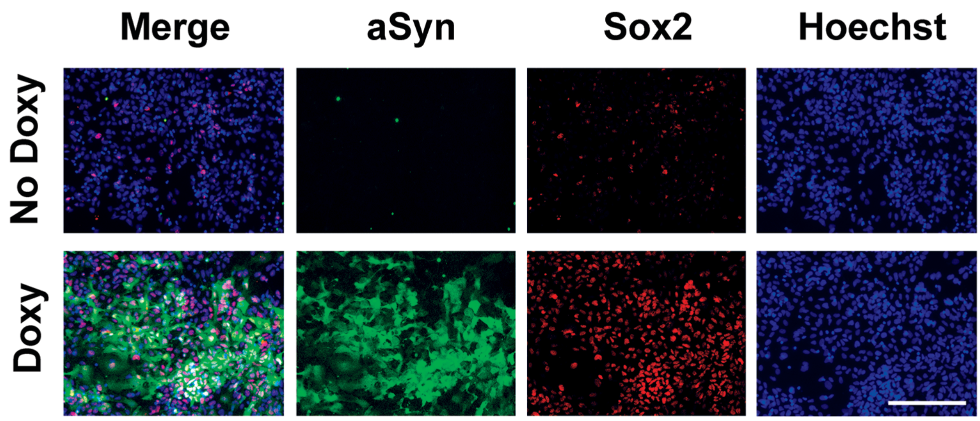


**Figure S3.** aSyn induction reduced neurogenesis in in AF22 Tet-On aSyn cells.

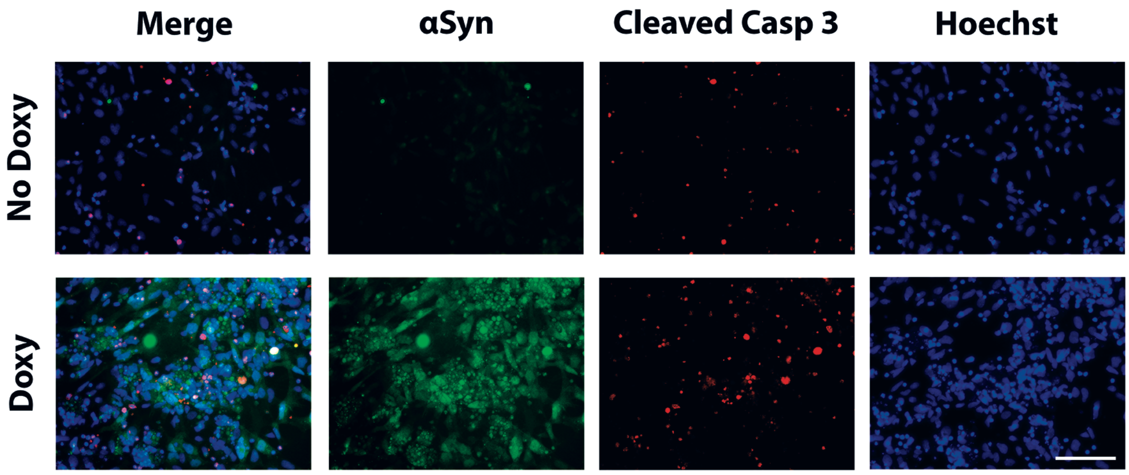


**Figure S4.** Acute aSyn induction impairs AF22 Tet-On aSyn cell-derived dopaminergic neurons viability.

| Antigen | Company | Dilution | Species |
|---|---|---|---|
| **α-synuclein** | Sigma | 1:1000-1:500 | Mouse |
| **β3-Tubulin** | Promega | 1:1000 | Mouse |
| **GFAP** | DAKO | 1:1000 | Mouse |
| **Sox2** | Millipore | 1:300 | Rabbit |
| **Phospho-Histone H3** | Chemicon/Millipore | 1:500 | Rabbit |
| **Cleaved Caspase 3** | CellSignalling Technology | 1:1000 | Rabbit |
| **Map2** | Millipore | 1:300 | Rabbit |
| **Nurr-1** | Santa Cruz Biotecnologies | 1:100 | Rabbit |
| **TH** | Sigma | 1:500 | Rabbit |
| **Alexa Fluor IgG anti-rabbit 568** | Molecular Probes | 1:500 | Goat |
| **Alexa Fluor IgG anti-rabbit 488** | Molecular Probes | 1:500 | Goat |
| **Immunostar anti-mouse HRP** | Biorad | 1:2000 | Goat |
| **Immunostar anti-rabbit HRP** | Biorad | 1:2000 | Goat |

**Table S1.** List of primary and secondary antibodies used in the study.
